# Uncovering the Environmental Conditions Required for *Phyllachora maydis* Infection and Tar Spot Development on Corn in the United States for Use as Predictive Models for Future Epidemics

**DOI:** 10.1101/2023.03.18.533264

**Authors:** Richard W. Webster, Camila Nicolli, Tom W. Allen, Mandy D. Bish, Kaitlyn Bissonette, Jill C. Check, Martin I. Chilvers, Maíra R. Duffeck, Nathan Kleczewski, Jane Marian Luis, Brian D. Mueller, Pierce A. Paul, Paul P. Price, Alison E. Robertson, Tiffanna J. Ross, Clarice Schmidt, Roger Schmidt, Teryl Schmidt, Sujoung Shim, Darcy E. P. Telenko, Kiersten Wise, Damon L. Smith

## Abstract

*Phyllachora maydis* is a fungal pathogen causing tar spot of corn (*Zea mays* L.), a new and emerging, yield-limiting disease in the United States. Since being first reported in Illinois and Indiana in 2015, *P. maydis* can now be found across much of the corn growing of the United States. Knowledge of the epidemiology of *P. maydis* is limited but could be useful in developing tar spot prediction tools. The research presented here aims to elucidate the environmental conditions necessary for the development of tar spot in the field and the creation of predictive models to anticipate future tar spot epidemics. Extended periods (30-day windowpanes) of moderate ambient temperature were most significant for explaining the development of tar spot. Shorter periods (14- to 21-day windowpanes) of moisture (relative humidity, dew point, number of hours with predicted leaf wetness) were negatively correlated with tar spot development. These weather variables were used to develop multiple logistic regression models, an ensembled model, and two machine learning models for the prediction of tar spot development. This work has improved the understanding of *P. maydis* epidemiology and provided the foundation for the development of a predictive tool for anticipating future tar spot epidemics.

## Introduction

Tar spot, caused by *Phyllachora maydis,* is an emergent disease on corn (*Zea mays* L.) that can lead to significant yield losses in the United States^1, 2^. First recorded infecting corn in Mexico as early as 1904^3^, *P. maydis* has since been reported throughout much of Latin America^4^. *Phyllachora maydis* had never been documented in the United States until 2015 when tar spot was observed in multiple fields in northern Indiana and Illinois^5^. Since its arrival in the United States, *P. maydis* has rapidly spread throughout the midwestern corn belt of the United States (U.S.). It has also been found in Florida, and Ontario, Canada^6^, along with confirmations in Georgia and Virginia^7, 8^. Under ideal environmental conditions, tar spot can cause severe epidemics. In 2018 alone, tar spot caused estimated yield losses of close to 5 million metric tons, equating to over 680 million USD of economic losses^2^.

Despite *P. maydis* being recognized as a pathogen of corn for over 100 years, there is still little understanding of its biology and epidemiology. As with other pathogens, environmental conditions greatly influence pathogen survival, dispersal, infection, and growth. *Phyllachora maydi*s overwinters in the U.S. corn production regions, indicating the pathogen can survive on corn residue, and consequently serves as the inoculum source for at least the next season’s crop^9, 10^. Monthly temperatures of 17°C – 22°C, relative humidity greater than 75%, leaf wetness of seven hours per night and 10-20 foggy days per month were reported as the optimal conditions for tar spot development^11^. Under controlled environments and optimal conditions, inoculation assays demonstrated a latent period of only fifteen days, and sporulation occurring approximately twenty days post-inoculation^12^.

As *P. maydis* continues to establish itself across several states in the U.S., an integrated management approach to mitigate the yield losses is needed. Partial genetic resistance for tar spot has been identified in corn germplasm^13, 14^, but many current commercial corn hybrids are considered highly susceptible. Fungicides are currently the most effective method for reducing tar spot development and yield losses, especially when two or three fungicide classes are used^15^.

In conjunction with the use of fungicides, predictive modeling has been an effective tool for guiding the optimal timing of fungicide applications. Predictive models have been developed for a multitude of varying pathosystems including Fusarium head blight of wheat primarily caused by *Fusarium graminearum* in the U.S.^16–19^, late blight of potato caused by *Phytophthora infestans*^20^, Sclerotinia stem rot of soybean caused by *Sclerotinia sclerotiorum*^21, 22^, and fire blight of apple and pear caused by *Erwinia amylovora*^23, 24^. Many of these models have been integrated into decision support systems, allowing farmers access to the predictive abilities of these models. One successful example is Sporecaster (https://ipcm.wisc.edu/apps/sporecaster/), a decision support system for the prediction of Sclerotinia stem rot of soybean which is publicly available to farmers on smartphones^22^.

Historically, many predictive models have been developed using either linear or logistic regression models^19–22, 24, 25^, but more recently predictive model development has shifted towards machine learning based analyses^26, 27^. One commonly used machine learning algorithm is a random forest (RF), which is an ensemble learning method for regressions^28^. The RF framework utilizes an aggregation of many decision trees allowing improved precision by reducing the amount of variance relative to single decision trees. However, RFs are not capable of being easily interpreted and overfitting can often occur. Another common machine learning algorithm is an artificial neural network^29^ (ANN), which resembles the interconnectedness and signaling of biological neurons. ANNs are made up of an input layer, consisting of either a single or multiple hidden layers, and an output layer. Within this network there are multiple nodes which are connected to many additional nodes, each of which carries their own associated weight and threshold. If the output from a single node meets a designated threshold, the node is triggered and sends data to the next layer of nodes. If a node does not meet the designated threshold, the node does not send data to the next layer. The downsides to using ANNs are the high level of complexity making it difficult to interpret the models, the considerable amount of computational power required to run these models, and the potential for overfitting. However, machine learning algorithms have been demonstrated to be highly effective at improving predictive capabilities due to their ability to model very complex and non-linear relationships^27, 30^.

The surge of new and/or re-emerging plant diseases represents one of the biggest challenges to food production in modern agriculture. As global climate change leads to instability of temperatures and changing precipitation patterns, the need to create greater resilience in our crop production systems has become crucial^31^. There are several knowledge gaps regarding tar spot development on corn. The tar spot cycle is not fully understood, specifically, the incubation and latent periods have not been clearly established for *P. maydis* in production settings, and information on pathogen dispersal is limited. Knowledge of these processes is critical in understanding the polycyclic nature of tar spot epidemics. Therefore, the goals of the current study were to discern the environmental variables that are most important for the development of tar spot, and to develop statistical models for the prediction of future tar spot epidemics in the U.S. that would maximize the precision of in-season management decisions.

## Results

### Development of Training and Testing Datasets

From this study, a dataset was compiled with 588 observations across the Midwest region of the U.S. including a binary response variable for the increase in *P. maydis* stroma between two consecutive rating dates. Of these 588 observations, 179 observations were taken from small-plot research trials between 2018 and 2022, and the additional 409 observations were taken from production fields between 2020 and 2022 (Fig. 1). From the combined 588 observations, designated training and testing data sets were created using a 70:30 split by randomly sampling from the full data set (small-plot and commercial fields combined) with replacement in which 70% of the observations were placed in the training data set and 30% of the observations were placed in the testing data set. The training data set included 96 observations where *P. maydis* developed or increased in severity and 310 observations in which *P. maydis* did not develop or increase in severity from the previous date. The testing data set included 36 observations in which *P. maydis* did increase in severity and 146 observations in which *P. maydis* did not develop or increase in severity. After the development of these two datasets, the training dataset was used for assessment of weather parameters and model development, while the testing data set was used for validation of the developed models from the training data set.

**Figure 1.**
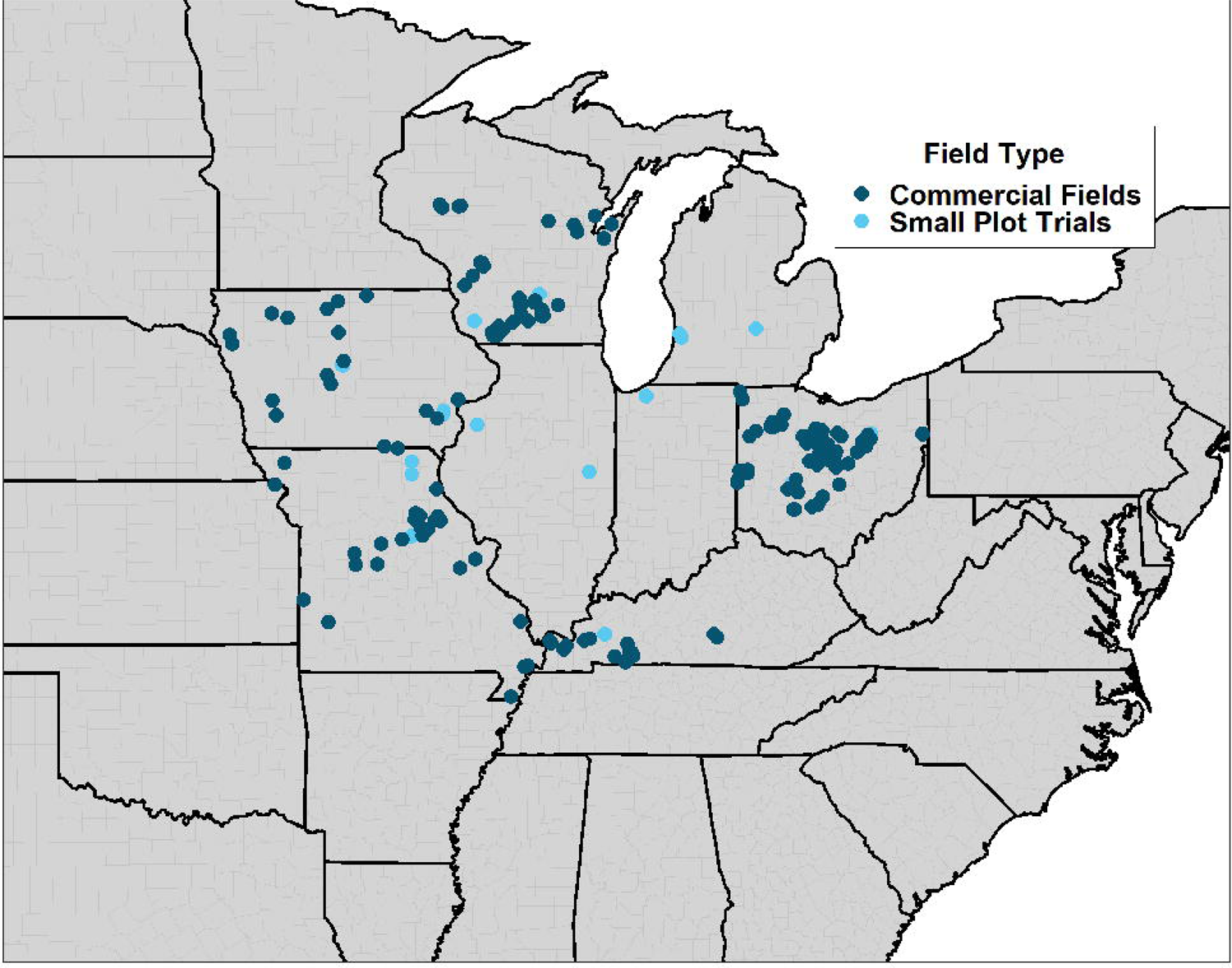
Map of all field locations where data were recorded and included in this study.

### Assessment of Weather Parameters

Multiple weather variables from the IBM historical weather data service were examined in this study across three levels of moving averages (windowpanes), 30-day (Fig. 2A), 21-day (Fig. 2B), and 14-day (Fig. 2C). By evaluating Pearson correlation coefficients of these moving averages in relation to the delta response variable, the strongest correlations were detected for the 30-day moving averages of the daily minimum ambient temperature and the daily mean ambient temperature with coefficients of −0.39 and −0.38, respectively (Fig. 2A, Suppl. Table 1). Within the 21-day moving averages, the two variables with the strongest correlations to *P. maydis* development or increase were the daily minimum dew point and the daily minimum temperature with coefficients of −0.36 and −0.35, respectively (Fig. 2B, Suppl. Table 1). Overall, there were eight 30-day moving average variables, fifteen 21-day moving average variables, and sixteen 14-day moving average variables significantly correlated with *P. maydis* development or increase in severity (Fig. 2, Suppl. Table 1).

**Figure 2.**
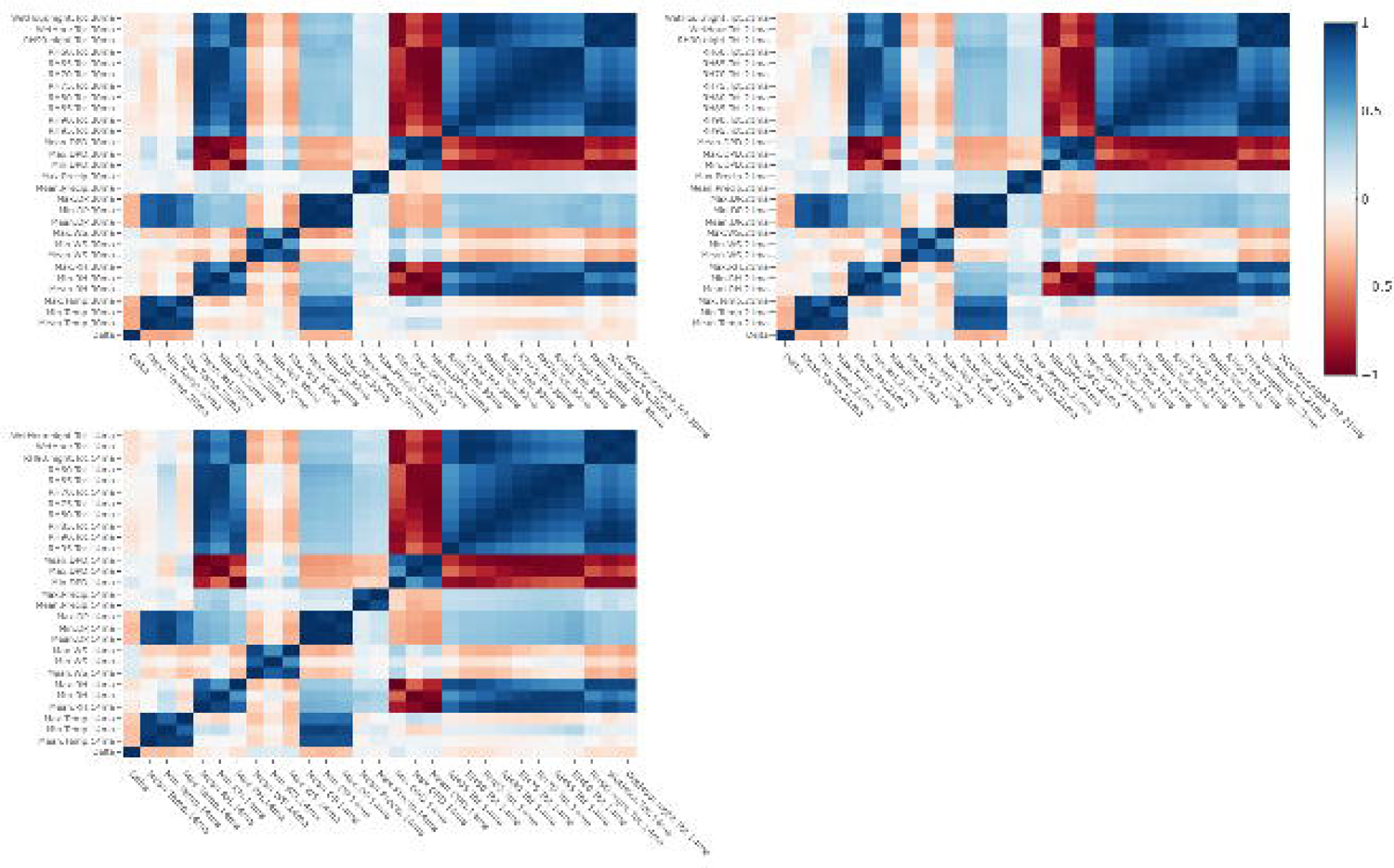
Pearson correlation matrix of the binary delta variable and 30-day moving averages (A), 21-day moving averages (B), and 14-day moving averages (C). Hyperlink can be used to view the interactive figure. https://chart-studio.plotly.com/~richard.webster/1/#plot.

As previously proposed by Hock et al.^11^, relative humidity (RH) levels are highly important in explaining *P. maydis* presence or increase in severity, especially at mean RH levels of 75% or greater. To investigate the impact of RH on *P. maydis*, we evaluated multiple 30-day moving averages of daily total hours of RH levels that ranged from 60% to 95% at 5% increments. Daily total hours of RH greater than 90% was significantly correlated with *P. maydis* development or severity increase for all three levels of moving averages (Fig. 2, Suppl. Table 1). Furthermore, daily total hours of RH greater than 85% was also significantly correlated with *P. maydis* development or severity increase for the 21-day and 14-day moving averages. Since these results suggested the importance of the 90% RH threshold, we also investigated the correlation of nighttime total hours of RH greater than 90% between 8 pm and 6 am. Nighttime total hours of RH greater than 90% was more highly correlated in all three levels of moving averages than the originally assessed values (Suppl. Table 1). However, the majority of correlations for the discussed RH variables were negatively correlated with *P. maydis* development or severity increase (Fig. 2, Suppl. Table 1). Additionally, a daily total wetness hour parameter was assessed serving as a proxy for the presence of leaf wetness. Like RH greater than 90% at night, the wetness hour parameter was evaluated as two distinct parameters for the total daily hours with predicted wetness and the total nighttime hours with predicted wetness. Both wetness hour parameters were not significantly correlated with *P. maydis* development or severity increase in the 30-day moving averages but were significant for both the 21-day and 14-day moving averages. The total nighttime wetness hours parameter was most highly correlated at the 14-day moving average with a correlation coefficient of −0.17 (*P* = 0.001, Fig. 2, Suppl. Table 1).

All assessed weather variables were used to create single variable logistic regression (LR) models for explaining *P. maydis* development and severity increase. These models were then evaluated by comparing Akaike information criterion (AIC) values, C-statistic values, and Hosmer-Lemeshow goodness-of-fit test *P*-values. From these evaluations, the two best fitting models were the models using 30-day moving averages of either the daily minimum temperature or the daily mean temperature (Fig. 3, Suppl. Table 2). When these two parameters were examined on the predicted risk probability, the inflection points observed for the daily minimum temperature was 15.4℃ and the daily mean temperature was 20.5℃ (Fig. 3).

**Figure 3.**
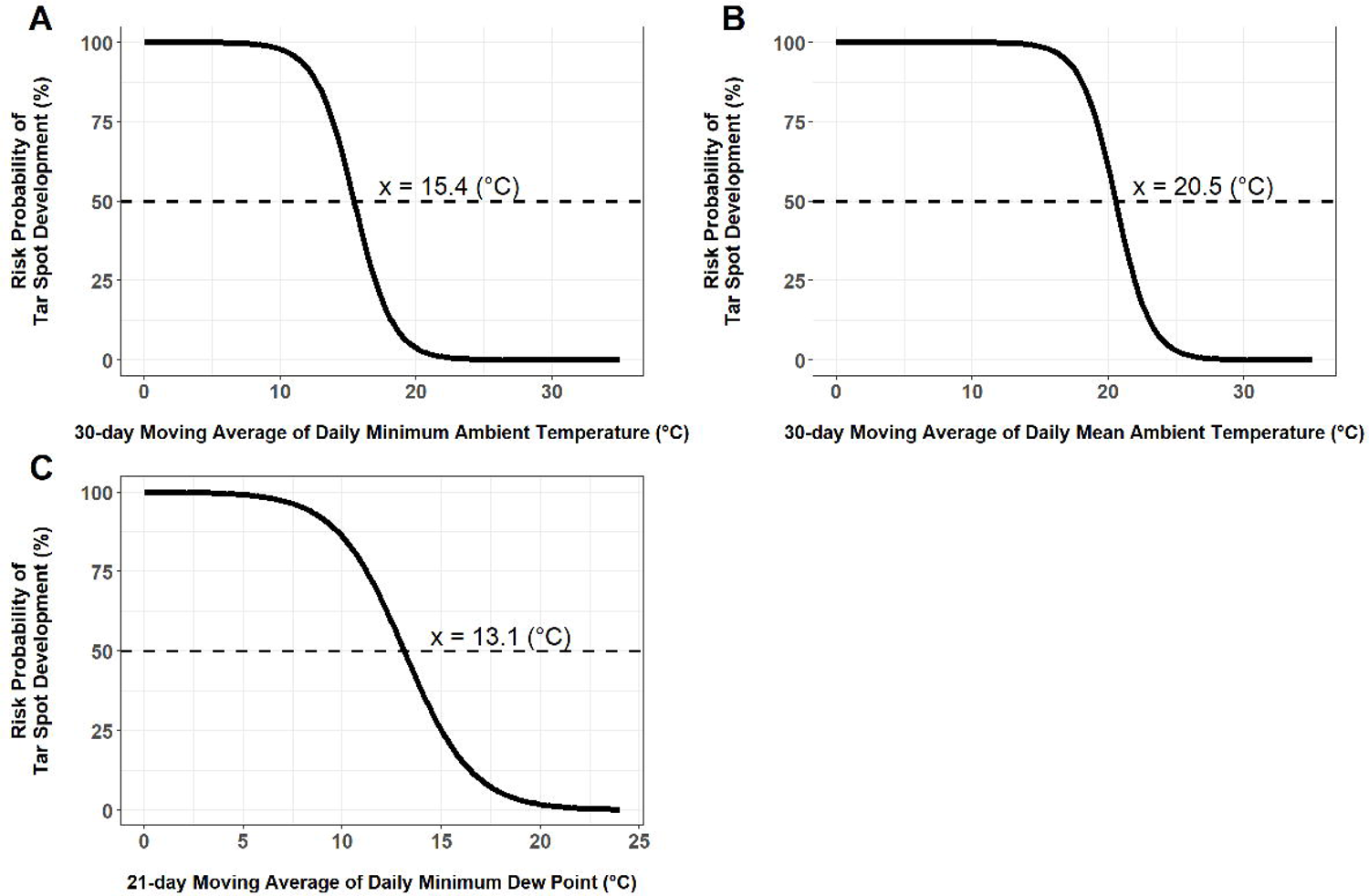
Logistic regression models developed for predicting the development of tar spot caused by *Phyllachora maydis.* These models predict the risk probability (%) of tar spot developing in relationship with (A) 30-day moving average of daily minimum ambient temperature (°C), (B) 30-day moving average of daily mean ambient temperature (°C), or (C) 21-day moving average of daily minimum dew point (°C).

Many of the moisture related parameters were best fitting when using either the 14-day or 21-day moving averages, compared to the 30-day moving average. For example, the daily minimum DP was observed to be best fitting when using the 21-day moving averages compared to the 14- and 30-day moving averages (Suppl. Table 2). From predicted risk probabilities as a result of the 21-day moving average of the daily minimum DP, an inflection point of 13.1℃ (Fig. 3). In addition, the nighttime total hours of RH greater than 90% and the total night time hours with predicted wetness parameters were best fitting when using the 14-day moving averages. This suggests the importance of reduced moisture in the 14 to 21 days prior to *P. maydis* development or increase in severity (Suppl. Table 2).

### Development of Predictive Models

With many single variable models developed, multi-variable models were then developed using the results of the previous assessments. Since the 30-day moving average of the daily minimum temperature and the daily mean temperature were the two most influential variables (Fig. 2, Suppl. Table 1 and 2), these two variables were examined more closely. Daily mean temperature was consistently more influential than the daily minimum temperature, and thus this variable was included in all subsequent models, which included many moisture variables. Eight models were chosen based on their input variables and favorable statistics reported above. Four of these models used the 30-day moving averages and included the combination of the daily mean temperature with either the daily total hours of RH greater than 90%, daily total wetness hours, daily minimum dew point depression (DPD), or the daily maximum RH (Suppl. Table 2). After these models were developed, the combination of different moving averages of weather parameters were explored due to the difference in influence as presented by the correlation coefficients and the single variable LRs (Suppl. Table 1 and 2). Four models were identified which all included the 30-day moving average of the daily mean temperature in addition to the 21- and 14-day moving averages of either the daily total hours with RH greater than 90%, the daily total nighttime hours with RH greater than 90%, the daily total wetness hours, or the daily total of nighttime wetness hours.

The corresponding eight LR models (LR1 – LR8) were selected to be validated using the previously established testing dataset. The linearized logistic models for these eight LRs are defined as:

1. Logit_LR1_ = 21.92522 − 0.97199(30 day moving average mean temperature) − 0.25014(30 day moving average of daily total of hours with RH > 90)
2. Logit_LR2_= 22.6108 − 0.9880(30 day moving average mean temperature) − 6.0357(30 day moving average of daily mean wetness hours)
3. Logit_LR3_ = 17.7869 − 0.8964(30 day moving average mean temperature) + 0.8157(30 day moving average of daily minimum dew point depression)
4. Logit_LR4_ = 32.06987 − 0.89471(30 day moving average mean temperature) − 0.14373(30 day moving average of daily maximum relative humidity)
5. Logit_LR5_ = 21.21170 − 0.94178(30 day moving average mean temperature) − 0.23661(21 day moving average of hours with RH > 90)
6. Logit_LR6_ = 20.35950 − 0.91093(30 day moving average mean temperature) − 0.29240(14 day moving average of daily total nighttime hours with RH > 90)
7. Logit_LR7_ = 22.18844 − 0.96662(30 day moving average mean temperature) − 0.25134(21 day moving average of daily total of wetness hours)
8. Logit_LR8_ = 21.66220 − 0.94504(30 day moving average mean temperature) − 0.34001(14 day moving average of daily total nighttime wetness hours)

The eight models were then assessed based on multiple model quality characteristics including accuracy (%), kappa value, type I error (%), type II error (%), precision (%), and recall (%) when using a 35% risk probability threshold for the prediction of *P. maydis* development or severity increase. From these evaluations, LR6 had the greatest accuracy (86.8%), greatest kappa value (0.59), greatest recall (69.4%), and the lowest type II error rate (30.6%, Table 1). Furthermore, LR4 had some of the best values for accuracy (86.3%), kappa (0.56), type I error rate (8.2%), and precision (65.7%, Table 1). From these results, a multi-model ensemble was created using LR4 and LR6 (Fig. 4) to more robustly predict the development of *P. maydis* or increase in its severity. The corresponding multi-model ensemble improved the accuracy (87.4%), kappa value (0.61), and precision (67.6%) while maintaining low type I error rate (8.2%), low type II error rate (30.6%), and high recall (69.4%, Table 1).

**Figure 4.**
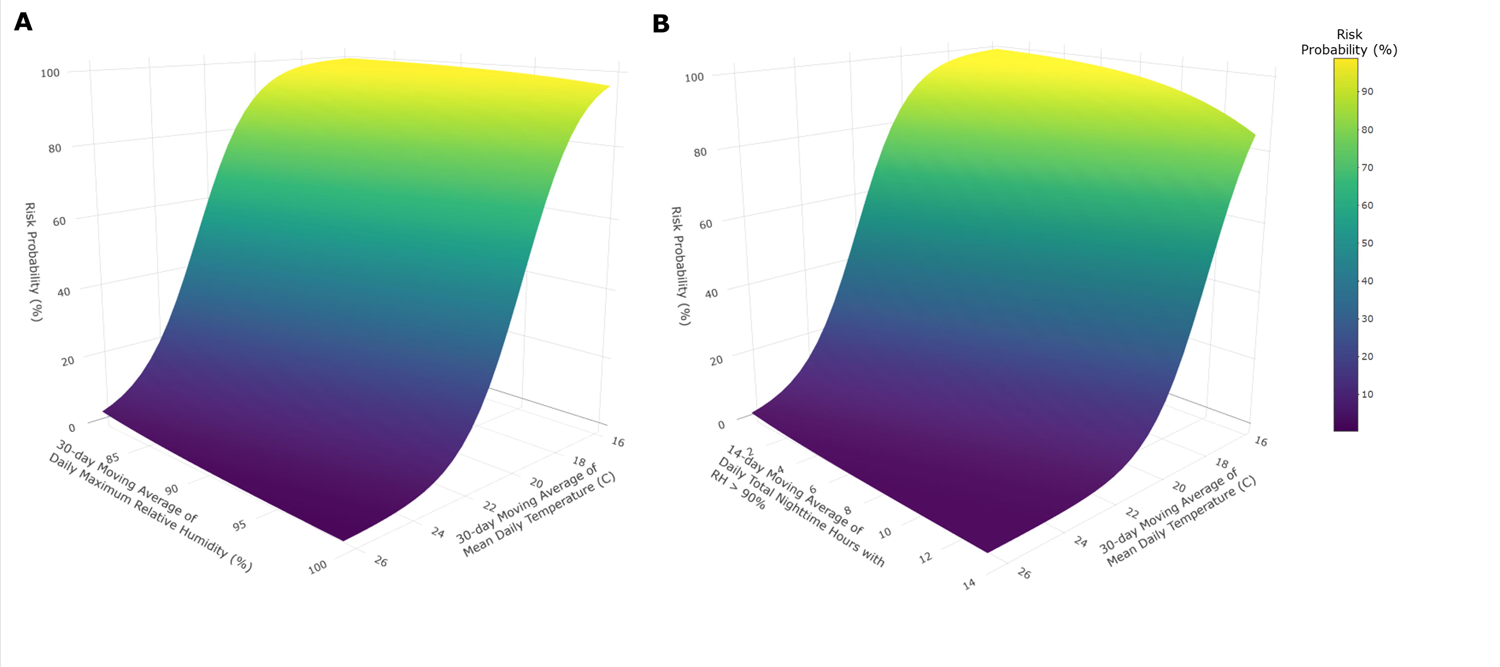
Two dimensional surfaces of logistic regression models developed for predicting the development of tar spot caused by *Phyllachora maydis.* (A) Logistic regression 4: Risk probability (%) of tar spot with 30-day moving average of daily mean ambient temperature (°C) and either 30-day moving average of daily maximum relative humidity or (B) Logistic regression 6: 14-day moving average of total nighttime hours with relative humidity > 90 %.

**Table 1.**
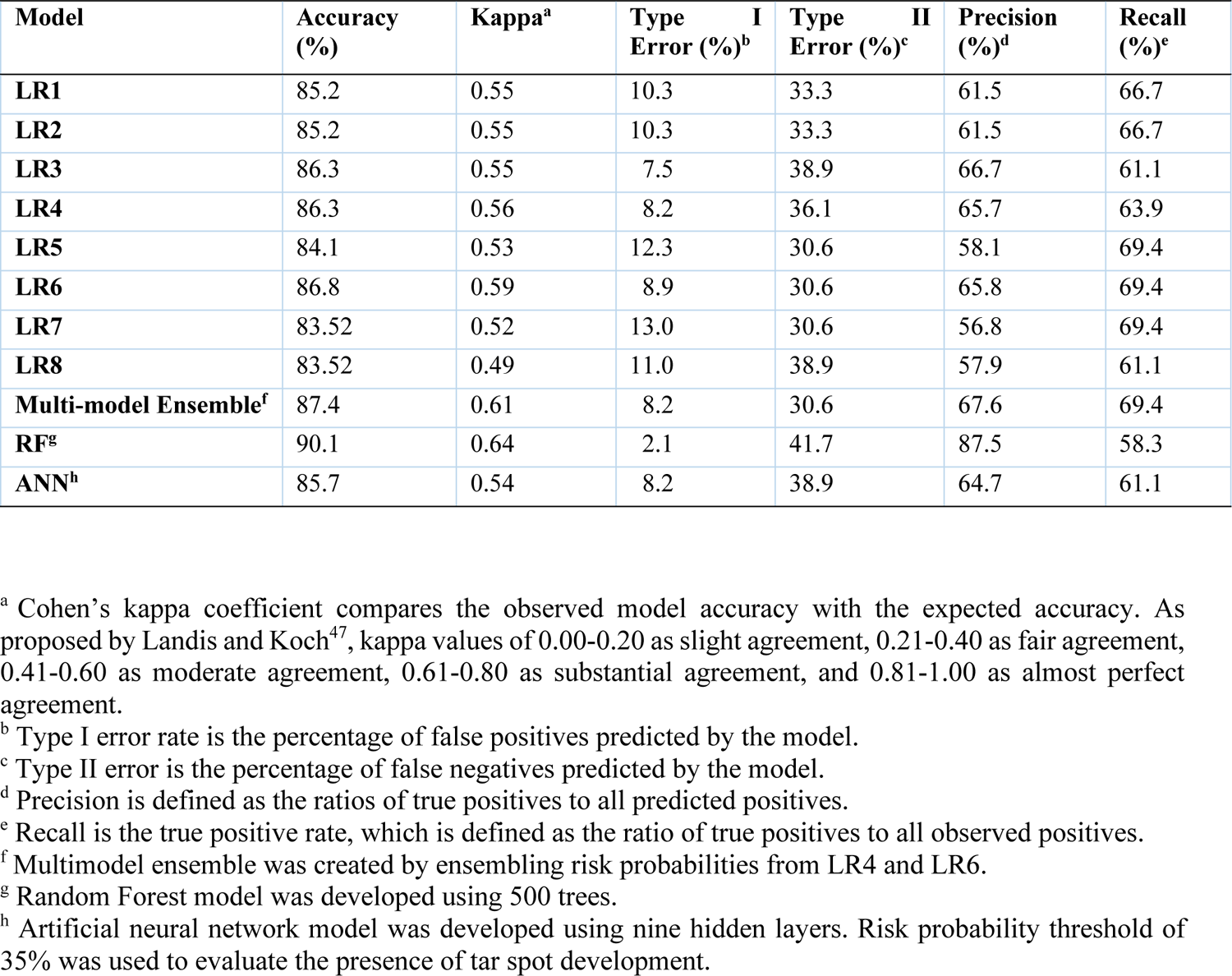
Model evaluation metrics for eight logistic regression models (LR1-LR8), a multi-model ensemble, a random forest model (RF), and an artificial neural network model (ANN) for predicting the development of tar spot (tar spot) on corn between 2018 and 2022 (n = 182).

Furthermore, two machine learning algorithms were developed for the prediction of *P. maydis* development and increase in severity including a RF model using 500 trees and an ANN model using nine hidden layers. These two machine learning algorithms were assessed similarly to the previous LRs and the multi-model ensemble. The RF consistently outperformed all other models for every metric except for recall, in which it had the lowest observed value (58.3%, Table 1). The corresponding RF had an observed accuracy of 90.1%. The ANN resulted in a model accuracy of (85.7%), relatively high type II error rate (38.9%), and all other metrics were unremarkable (Table 1).

## Discussion

Through this study, a deeper epidemiological understanding of *P. maydis* has been uncovered. The corresponding research suggests the development and increase in *P. maydis* stroma under field conditions is primarily driven by extended periods of moderate ambient temperature and short periods of high relative humidity. Additionally, the development of multiple statistical models offers a tool in production systems to guide fungicide applications to help farmers maximize fungicide efficacy while balancing return on investment.

From our findings, the weather parameters with the strongest correlations to *P. maydis* development and severity increase included the 30-day moving averages of either daily minimum temperature or the daily mean temperature (Suppl. Table 1). These two weather parameters were also correlated with *P. maydis* development and severity increase from the 21-day and 14-day moving averages but were not as highly correlated as the 30-day moving averages (Suppl. Table 1). Specifically, moderately warm air temperatures appear to drive *P. maydis* development or severity increase, while excessively warm conditions, specifically mean temperatures greater than 20.5°C, considerably decreased the probability of *P. maydis* progression (Figure. 2B). This demonstrates there is a long-term influence of moderate ambient temperature that drives epidemiological processes within the tar spot cycle. These results confirm previous reports by Hock et al.^11^ that moderate temperatures (17-22℃) were one of the primary determinants of tar spot progress and severity. Hock et al.^11^ also reported that during warm seasons with mean ambient temperatures greater than 22℃, tar spot development was minimal. Moderate mean temperatures could be influencing multiple epidemiological processes within the disease cycle such as germination of initial inoculum, infection of the host, mycelial colonization within host tissue, or the development of the ascomata. Temperature has been well characterized to play an important role in all of these processes in many other fungal organisms^21^. Additional investigations on the effects of temperature on the development of *P. maydis* on corn are still needed to further elucidate this relationship.

In addition to temperature, moisture weather variables were consistently observed to influence tar spot development. Specifically, the 21-day moving average of the daily minimum dew point (DP) had the third greatest overall correlation with *P. maydis* development or severity increase (Fig. 2, Suppl. Table 1). Many additional moisture parameters were significantly correlated with *P. maydis* development or severity increase across all three levels of moving averages, such as the RH90 and the nighttime RH90 variables as has been previously reported (Fig. 2). Interestingly these moisture variables are negatively correlated in our work. Breunig et al.^12^ point out that in controlled-environment inoculations frequent misting was only required in the first five days after inoculation. After five days, misting had to be withheld to produce stroma in these controlled environments. Perhaps leaf wetness is required by the fungus for spore germination and leaf penetration, while excessive moisture later in the infection process can lead to washing or other detrimental events that don’t promote infection. Regardless, these moisture variables are clearly playing a role in the biological processes driving the development of *P. maydis*. Variables such as DP and RH are still dependent on temperature. Thus, the relationships presented here demonstrate the complexity that exists between the roles of ambient temperature and moisture on *P. maydis* development in the physical environment. As Hock et al.^11^ reported, RH levels greater than 75% were important for tar spot development. We examined the effect of different RH thresholds ranging from 60% up to 95%, and consistently determined the 90% RH threshold was most influential in explaining the development and increase in severity of *P. maydis*, with RH90 being significantly correlated with *P. maydis* stroma development in all three levels of moving averages (Suppl. Table 1) and resulting in the best fitting models across all RH thresholds (Suppl. Table 2). Our data are like those of Hock et al.^11^, in that RH was most important in predicting *P. maydis* development. Thus, the results presented here refine our understanding of the role RH plays in the epidemiology of tar spot in the U.S.

Another important objective of this study was to compare LR models to more modern machine learning algorithms. From our study, a RF machine learning algorithm resulted in one of the best models and had the greatest observed model accuracy of 90.1%. The ANN examined in this study did not result in model accuracy as high as several of the LR models developed here. Two LRs, LR4 and LR 6, were highly accurate. Accuracy was further improved by ensembling these two LR models, with an accuracy estimate of 87.4% while either improving or maintaining all of the additional model assessment characteristics (Table 1). Our analyses demonstrate that machine learning algorithms were slightly more accurate in predicting *P. maydis* development compared to LRs, but a multi-model ensemble using two of the LRs was still comparable in predicting *P. maydis* development while balancing all goodness-of-fit statistics. These results confirm previous studies on predicting plant diseases with a high degree of accuracy using different machine learning algorithms^27, 32, 33^. However, in some studies logistic regressions were reported to still be the most accurate at predicting plant diseases^34^.

Functionally, LR models may be more useful in actual delivery to farmers and can be easily programmed into smartphone application decision support systems (DSS) as has been previously demonstrated^22^. The models presented here have high levels of potential for improving the application timings of fungicides for managing tar spot. Tar spot may result in severe yield losses; thus, farmers often rely on multiple fungicide applications during the season, which can equate to high economic and environmental costs. The use of these DSS will guide farmers in optimizing fungicide application timing to protect the plant from the pathogen when it is most likely to cause disease. Furthermore, using these DSS can also eliminate unnecessary applications which benefit the farmers by limiting needless economic inputs and decreasing chemical inputs into the environment.

While these identified models are highly accurate at predicting *P. maydis* development, there is inherit error associated with any model. This error could be explained by variability in environmental conditions which could not be accounted for, the quantity of initial inoculum, the population structure of the pathogen within the field, or resistance levels among site-years. Additionally, there may have been interrater variability associated with disease ratings, especially since these data were collected from multi-state projects with numerous raters across multiple years. However, we minimized this error with the use of standard area diagrams (CPN) that help improve disease severity estimates^35^.

The current study sheds light on epidemiological processes that are driving the development of a newly emerged pathogen of corn capable of causing severe disruptions to agricultural production. The work presented here has also paved the way for the development of a DSS for tar spot. Work is underway to incorporate these models into the Tarspotter DSS (https://ipcm.wisc.edu/apps/tarspotter/) to further improve tar spot prediction and better inform farmers of risk due to plant disease.

## Material and Methods

### Field Trials

Small plot field trials were planted between 2018 and 2022 in the following states: Illinois, Iowa, Indiana, Kentucky, Michigan, Missouri, Ohio, and Wisconsin (Fig. 1). Locally adapted hybrids were used at each location. The use of all plant material in this study did not require any specific permissions or licenses. Trials at each location followed locally recommended management practices such as seeding rates, nitrogen fertilization, and herbicides with a small number of trials overhead irrigated. Field trials in 2018 and 2019 included the use of fungicide applications, but only the non-treated plot was considered for this study. No fungicide applications were made in trials conducted between 2020 and 2022. Commercial field sites were also assessed across the Midwest U.S. between 2021 and 2022. The conducted field trials were performed with permission from local commercial grower collaborators and were compliant with all institutional, national, and international guidelines and legislation. Additional field information is provided in Supplementary Table 3.

### Data Collection

*Phyllachora maydis* ratings started at the R1 growth stage (silking) and continued until the R5-R6 growth stage (dent to full maturity). The number of *P. maydis* severity ratings during this period ranged from two to seven depending on the site-year. *Phyllachora maydis* severity was rated by visually assessing the percentage of *P. maydis*-induced stroma on the ear leaf of five to ten plants per plot (sub-samples) using a standardized rating scale^36^, and all ratings were averaged across the entire plot. For each rating date in each site-year, all plot severity scores were averaged for a single severity score for that plot. The compiled database considered for developing the prediction modeling was the average *P. maydis* severity of the ear leaf for each assessment day. The severity data were aggregated into a single file. For each location, all ratings were aligned in sequential order by date. A binary delta variable was defined as the increase in severity of *P. maydis* stroma between two sequential rating dates, such that a delta value of 1 was given for any positive increase in *P. maydis* severity between two sequential dates. If no increase in *P. maydis* severity was observed, a delta value of 0 was given. Thus, delta values of 1 define *P. maydis* increase while delta values of 0 define no increase.

### Weather Data Collection

Site-specific weather data was collected using IBM historical weather services. Hourly average weather data was pulled from this service at a resolution of 4 km grids using GPS coordinates for each field location. The collected weather data included the hourly averages of ambient air temperature (AT), relative humidity (RH, %), wind speed (WS, m/s), dew point (DP), and precipitation (mm/hour). From these hourly weather data, dew point depression (DPD) values were calculated for each hour by taking the absolute value of the difference between the AT and DP. A binary wetness hour variable (WH) was calculated by defining a ‘1’ if the DPD was less than or equal to two, predicting the presence of free water on leaf surfaces, and a ‘0’ was defined if the DPD was greater than two^37^. Additionally, a binary nighttime wetness hour variable was calculated similarly to the previously described wetness hour variable but could only be considered true between the hours of 8 pm and 6 am. All other daytime hours were considered a ‘0’ value. A binary RH variable (RH95) was calculated by defining a ‘1’ if the RH was greater than or equal to 95%, and a ‘0’ was defined if the RH level was less than 95%. Additional binary RH variables (RH90, RH85, RH80, RH75, RH70, RH65, and RH60) were calculated similarly with RH thresholds of either 90%, 85%, 80%, 75%, 70%, 65%, or 60%. A binary nighttime RH90 variable was calculated by defining a ‘1’ if the RH was greater than 90% between the hours of 8 pm and 6 am, and if RH was less than 90% at night or during daytime hours all hours were defined as a ‘0’.

From these hourly weather values, daily mean, minimum, and maximum values were calculated for each of the following variables (AT, RH, WS, DP, and DPD). Daily mean and daily maximum precipitation rates were calculated. Daily totals for WH, nighttime WH, RH90, RH90, RH85, RH80, RH75, RH70, RH65, RH60, and nighttime RH90 were also calculated for each location. After all daily means, minimums, maximums, and totals were calculated, 30-day, 21-day, and 14-day moving averages (window-panes) were calculated for each of the weather variables using the rollmean() function from the ‘zoo’ package in R^38, 39^. “Window-paning” has been useful in modeling for Fusarium head blight for instance, allowing epidemiologists the ability to find and define specific time-frames for weather variables that are influential in plant disease development^40^. Finally, the previously established binary delta values were paired with the 30-day, 21-day, or 14-day moving averages of weather data for the second rating date.

### Correlation Analysis and Logistic Regression Model Development

First the total dataset was split to create training (n = 406) and testing (n = 182) datasets using bootstrapping with replacement. Correlation analyses were performed in R using the rcorr() function from the ‘Hmisc” package^41^. These analyses calculated the Pearson correlation coefficients for the delta values with respect to either 30-day, 21-day, or 14-day moving averages (windowpanes). Significant correlations were determined by a *P*-value of less than 0.05 (Suppl. Table 1). All LRs were developed with the delta variable as the response variable. Single variable LRs were created by using each of the 30-day, 21-day or 14-day moving averages for each of the weather parameters previously described. Additional multi-variable LRs were developed using a combination of these weather variables. All LRs were evaluated by Akaike information criterion (AIC) values, area under the receiver operating characteristics curve (C statistic) using the Cstat() function from the ‘DescTools’ package in R^42^, and tested by the Hosmer-Lemeshow goodness of fit test (HL test) using the hltest() function from the ‘glmtoolbox’ package in R^43^. Favorable models were determined as having the lowest AIC values, the highest C statistics, and a HL test *P*-value of greater than 0.05. From these assessments, eight LR models (LR1-LR8) were identified for further evaluation. Additionally, a multi-model ensemble was created by taking the daily average risk probability from the LR4 and LR6 models.

### Evaluation against Machine Learning Algorithms

To evaluate if the developed LR models were adequately predicting the progression of *P. maydis* on corn plants, the eight best-fitting LR models and ensemble model were compared against two different machine learning algorithms. These included random forests (RF) and artificial neural networks (ANN). From the training dataset, the delta response variable was examined to be explained by all predictor variables using the randomForest() function from the ‘randomForest’ package in R^44^ using a total of 500 trees and all other default hyperparameters were used. The subsequent RF model was then tested on the testing dataset to determine the accuracy of predicting the delta response variable. The training set was also used to create an ANN using the neuralnet() function from the ‘neuralnet’ package in R^45^ using nine hidden layers and all other hyperparameters were set to their default. This ANN was then used to evaluate the ability to predict the delta response variable from within the test dataset. Model fitness metrics compared to the testing data set for the eight LR models, the ensemble model, and the two machine learning models were evaluated for their accuracy (%), kappa values, type I error (%), type II error (%), precision (%), and recall (%). These metrics were evaluated for each model using the confusionMatrix() function from the ‘caret’ package in R^46^.

## Acknowledgements

This work was partially supported by the National Predictive Modeling Tool Initiative operating under the auspices of the USDA-ARS; the Wisconsin Corn Promotion Board; the Corn Marketing Program of Michigan; Project GREEEN - Michigan’s plant agriculture initiative; USDA National Institute of Food and Agriculture, Hatch Project #IND00162952; Foundation for Food Agricultural Research - Rapid Outcomes from Agricultural Research (FFAR-ROAR award # 0000000017) grant with matching funds provided by Pioneer, the National Corn Growers Association; The Illinois Corn Growers Association; Indiana Corn Marketing Council; Purdue University, and Hatch Project IOWN03908.

## Data Availability

Disease data are provided at https://github.com/rwwebster/Tar-Spot-Modelling At the time of submission, the authors have not yet made this data publicly available. However, the authors have included the raw data as an attached file titled “Disease Data.xlsx” for transparency during the review process. Data will be made publicly available upon full acceptance of the manuscript.

## Additional Information

The authors thank J. Ravellette and Steven Brand at Purdue University, John Boyse, Adam Byrne and William Widdicombe at Michigan State University, John Shriver and Cody Schneider at Iowa State University, Nolan Anderson, Jesse Gray, and Sean Wood at the University of Kentucky, and Dan Sjarpe at the University of Missouri for assistance with the field trial maintenance.

