## Supplemental Table Legends for "Uncovering the Environmental Conditions Required for *Phyllachora maydis* Infection and Tar Spot Development on Corn in the United States for Use as Predictive Models for Future Epidemics"

### **Supplementary Material**

Supplementary Table 1. Pearson correlation coefficient and significance p-value for 30-day moving average, 21-day moving average and 14-day moving average.

Supplementary Table 2. Weather parameters in logistic regression models for 30-day moving averages, 21-day moving averages, 14-day moving averages, and combined moving averages.

Supplementary Table 3. All information for small-plot trials and commercial fields planted in 2018 to 2022 in the following states: Illinois, Iowa, Indiana, Kentucky, Michigan, Missouri, Ohio, and Wisconsin in the United States.
